# Reward enhances online participants’ engagement with a demanding auditory task

**DOI:** 10.1101/2021.02.09.430467

**Authors:** Roberta Bianco, Gordon Mills, Mathilde de Kerangal, Stuart Rosen, Maria Chait

## Abstract

Online recruitment platforms are increasingly utilized for experimental research. Crowdsourcing is associated with numerous benefits but also notable constraints, including lack of control over participants’ environment and engagement. In the context of auditory experiments, these limitations may be particularly detrimental to threshold-based tasks that require effortful listening. Here, we ask whether incorporating a performance-based monetary bonus will improve speech reception performance of online participants. In two experiments, participants performed an adaptive matrix-type speech-in-noise task (where listeners select two key words out of closed sets). In Experiment 1, our results revealed worse performance in online (N = 49) compared with in-lab (N = 81) groups. Specifically, relative to the in-lab cohort, significantly fewer participants in the online group achieved very low (< -17dB) thresholds. In Experiment 2 (N = 200), we show that a monetary reward improved listeners’ threshold to levels similar to those observed in the lab setting. Overall the results suggest that providing a small performance-based bonus increases participants’ task-engagement, facilitating a more accurate estimation of auditory ability under challenging listening conditions.

## Introduction

There is a growing interest in remote testing, both in the context of basic research (Anwyl-Irvine, Dalmaijer, Hodges, & Evershed, 2020; Backx, Skirrow, Dente, Barnett, & Cormack, 2020; Hartshorne, de Leeuw, Goodman, Jennings, & O’Donnell, 2019; Lelo de Larrea-Mancera et al., 2020; Shapiro, Norris, Wilbur, Brungart, & Clavier, 2020) and clinical screening (De Wet & Clark, 2019; De Wet, De Sousa, Smits, & Moore, 2019; Paglialonga et al., 2020; Sevier, Choi, & Hughes, 2019; Shafiro et al., 2020; Sheikh Rashid, Dreschler, & de Laat, 2017; Watson, Kidd, Miller, Smits, & Humes, 2012). The ability to conduct experiments online facilitates rapid data acquisition and provides access to a larger and more diverse subject pool than that available for lab-based investigations (Casey, Chandler, Levine, Proctor, & Strolovitch, 2017). However, in contrast to the lab setting, online experiments are associated with a lack of control over participants’ equipment, environment and engagement (Chandler & Paolacci, 2017; Clifford & Jerit, 2014). These limitations may be particularly detrimental to auditory assessments which often rely on highly controlled stimulus delivery and necessitate focused engagement from the participant (e.g., Harrison & Müllensiefen, 2018).

Tasks that require effortful listening (e.g. when trying to estimate performance at threshold, or the just noticeable difference in a particular acoustic feature) may be particularly susceptible to issues related to task-engagement (including attention, motivation and commitment). In laboratories or clinics, engagement is controlled by creating a ‘sterile environment’ which isolates the participants from potential sources of distraction (e.g., their mobile phone, software notifications, doorbell, housemates, etc.). Compliance and motivation are promoted through face-to-face interaction with the experimenter (Guéguen & Pascual, 2000; Karakostas & Zizzo, 2016). To understand how these factors affect data obtained from online participants, in this series of experiments we investigated how performance on one version of widely used auditory speech-in-noise perception tasks differs between in-lab and online settings and whether monetary reward may be used as a means to encourage participant engagement.

We used an adaptive speech-in-noise task based on target materials similar to the Coordinate Response Measure (CRM) corpus of Bolia, Nelson, Ericson, & Simpson (2000). The CRM measures the ability to identify two key words (colour and number words) in a spoken target sentence always cued by a so called ‘call sign’. Participants are instructed to attend to the target sentence while ignoring a masker. The CRM is a powerful test of listening in complex environments because of its sensitivity to small intelligibility changes in highly noisy backgrounds, its applicability to testing with different maskers and its relative independence from semantic/syntactic cues (Brungart, 2001; Eddins & Liu, 2012; Humes, Kidd, & Fogerty, 2017). Accumulating work demonstrates that speech reception thresholds (SRT) estimated with an adaptive CRM task correlate with audiometric thresholds and with age (de Kerangal, Vickers, & Chait, 2020; Schoof & Rosen, 2014; Venezia, Leek, & Lindeman, 2020), rendering it a potentially efficient proxy of hearing ability (Semeraro, Rowan, van Besouw, & Allsopp, 2017). An additional advantage is that the task relies on manipulating the relative intensity of the target and the masker and performance is largely independent of overall level over a reasonable range. Outcomes are therefore less affected by calibration of equipment compared with other tasks that rely on absolute sound level. These considerations make the CRM, as well as other similar speech-in-noise tasks (De Sousa, Smits, Moore, Myburgh, & Swanepoel, 2020), particularly attractive for estimating auditory function in online settings.

We firstly asked whether performance among young listeners recruited “blindly” online is consistent with that observed in the highly controlled laboratory setting. Results suggested poorer performance by online listeners. We hypothesized that reduced performance in the online, compared with the in-lab, sample may reflect a lack of task engagement or motivation among the online cohort. Therefore, building on existing evidence that monetary reward can improve performance in tasks that involve executive or perceptual functions (Libera & Chelazzi, 2006; Shen & Chun, 2011), we asked whether incorporating a performance-based monetary bonus could improve speech reception performance in the online group. Our results revealed that a monetary bonus improved listeners’ thresholds to levels similar to those observed in the lab setting. Overall the results suggest that providing a small performance-based bonus may increase participant task engagement (i.e., the readiness to exert effort and/or allocate sufficient attention to the task) facilitating a more accurate estimation of auditory ability.

## Experiment 1

### Methods

#### Participants

Two participant groups ranging in age between 25 and 32 years were tested. An **in-lab group** (data pooled from de Kerangal, Vickers, & Chait, 2020 and an additional unpublished study) comprised 81 participants (59 females, mean age 25 ± 3 years) who completed the task as part of a test battery. An age matched **online group** of 49 participants (35 females, mean age 26 ± 3 years) was recruited and compensated via the Prolific crowdsourcing platform. All listeners were young, native speakers of British English, and reported no known hearing problems. Experimental procedures were approved by the research ethics committee of University College London and informed consent was obtained from each participant.

#### Stimuli and procedure

A speech recognition threshold (SRT) for each participant was obtained using target sentences introduced by Messaoud-Galusi, Hazan, & Rosen (2011) – the Children’s Coordinate Response measure (CCRM) which are a modified version of the CRM corpus described by (Bolia et al., 2000). The modifications were made in order to be able to embed the materials in the task as a straightforward command, and using call signs (here the animal name) that would be more appropriate for use with children, without precluding the use of the material in adults, nor changing the essential properties of the corpus. Note that the CCRM as used here is likely to be at least as difficult as the original CRM (both requiring the identification of a colour and a number) but here there are 6 colours rather than 4. On each trial, participants heard a target sentence of the form “show the dog where the [colour] [number] is.”. The number was a digit from 1 to 9, excluding the number 7 (due to its bisyllabic phonetic structure, which would make it easier to identify). The colours were black, white, pink, blue, green, or red. Thus, there were a total of 48 combinations (six colours times eight numbers). Participants were instructed to press on the correct combination of colour and number on a visual interface showing an image of a dog and a list of the digits in the different colours.

The target sentences were spoken by a single female native speaker of Standard Southern British English that was presented simultaneously with a 2 male-speaker babble that the participants were instructed to ignore. Each talker in the babble was recorded reading two 5- to 6-sentence passages which were concatenated together once passages were edited to delete pauses of more than 100 ms. The two talkers were then digitally mixed together at equal levels, with random sections of the appropriate duration from this 30 s long masker chosen for each trial.

The overall amplitude of the mixture (target speaker + babble background) was kept fixed, with only the ratio between the target and masker changing on each trial. The signal to noise ratio (SNR) between the babble and the target speaker was initially set to 20dB and was adjusted using a one-up one-down adaptive procedure, tracking the 50% correct threshold (Levitt, 1971). Initial steps were of 9dB, decreasing by 2dB following each reversal then stabilizing at a final step size of 3dB. The procedure terminated after 7 reversals. The SRT was calculated as the mean of the SNRs in the last 4 reversals. Each participant performed the test in 4 consecutive runs of approximately 2 minutes each; the mean over the SNRs collected in the last 3 runs was used for the analyses. The task took approximately 10 minutes.

The in-lab test was conducted in a double-walled sound-proof booth (IAC, Winchester). The task was implemented in MATLAB using a calibrated sound delivery system. Sounds were presented with a Roland Tri-capture 24-bit 96 kHz soundcard over headphones (Sennheiser HD 595) at a comfortable listening level (~60-70 dB SPL), self-adjusted by each participant during the training block.

For online testing, the task was implemented in javascript and the Gorilla Experiment Builder platform (www.gorilla.sc) was used to host the experiment (Anwyl-Irvine et al., 2020). Participants were recruited and pre-screened by the Prolific platform. Otherwise, the same stimuli and test heuristics were used as in the in-lab settings. To make sure that participants were wearing headphones, they were screened for headphone use with the approach introduced by Milne et al (2020). A strict version of the screening task, demonstrated to yield a 7% false positive rate, was used. The CCRM task began with a volume calibration to make sure that stimuli were presented at an appropriate level. A target sentence without a masker was used for this purpose. Participants were instructed to play the sound and adjust the volume to as high a level as possible without it being uncomfortable.

At the end of the experiment, participants completed a short questionnaire about their listening environment and equipment. We encouraged honest reports by stressing that “your answers will not affect your payment but will help us to get the best quality data”. In particular, participants were asked about how much background noise they experienced during the experiment (0 = not at all, 10 = a lot). This measure was used as a potential exclusion criterion to make sure that group differences in performance were not explained by mere differences in environmental noise. The task was piloted to take about 15 minutes. We thus set the base-pay rate to £ 2, corresponding to an hourly wage of £ 8.

We have made our implementation openly available and ready for use via Gorilla (https://gorilla.sc/openmaterials/171870).

#### Statistical analysis

We used the 2 sample Kolmogorov-Smirnov test to ascertain the existence of a statistically significant difference between the (unknown) distributions of the two groups of interest. Analyses were conducted in the R environment Version 0.99.320.

## Results

Figure 1 shows the probability density function (panel A) and the cumulative distribution function (panel B) of the SRT obtained from the in-lab (mean SRT = −16.2 dB, SD = 2.08) and online groups (mean SRT = −15.1 dB, SD = 2.21; mean difference in-lab – online = −1.1 dB). A KS test indicated a significant difference between the two distributions (D = .347, p = .001). The maximal difference occurred at −16.9 dB, which was reached by 47% of the in-lab group and only by the 12% of the online group. Despite the low level of background noise reported by the online sample (1.77 ± 2.51 from a range of 0-10), we repeated the analysis by excluding those participants who reported a high level of noise (>= 5 final sample N = 42). The difference between groups was unaltered (D = .350, p = .002).

**Figure 1.**
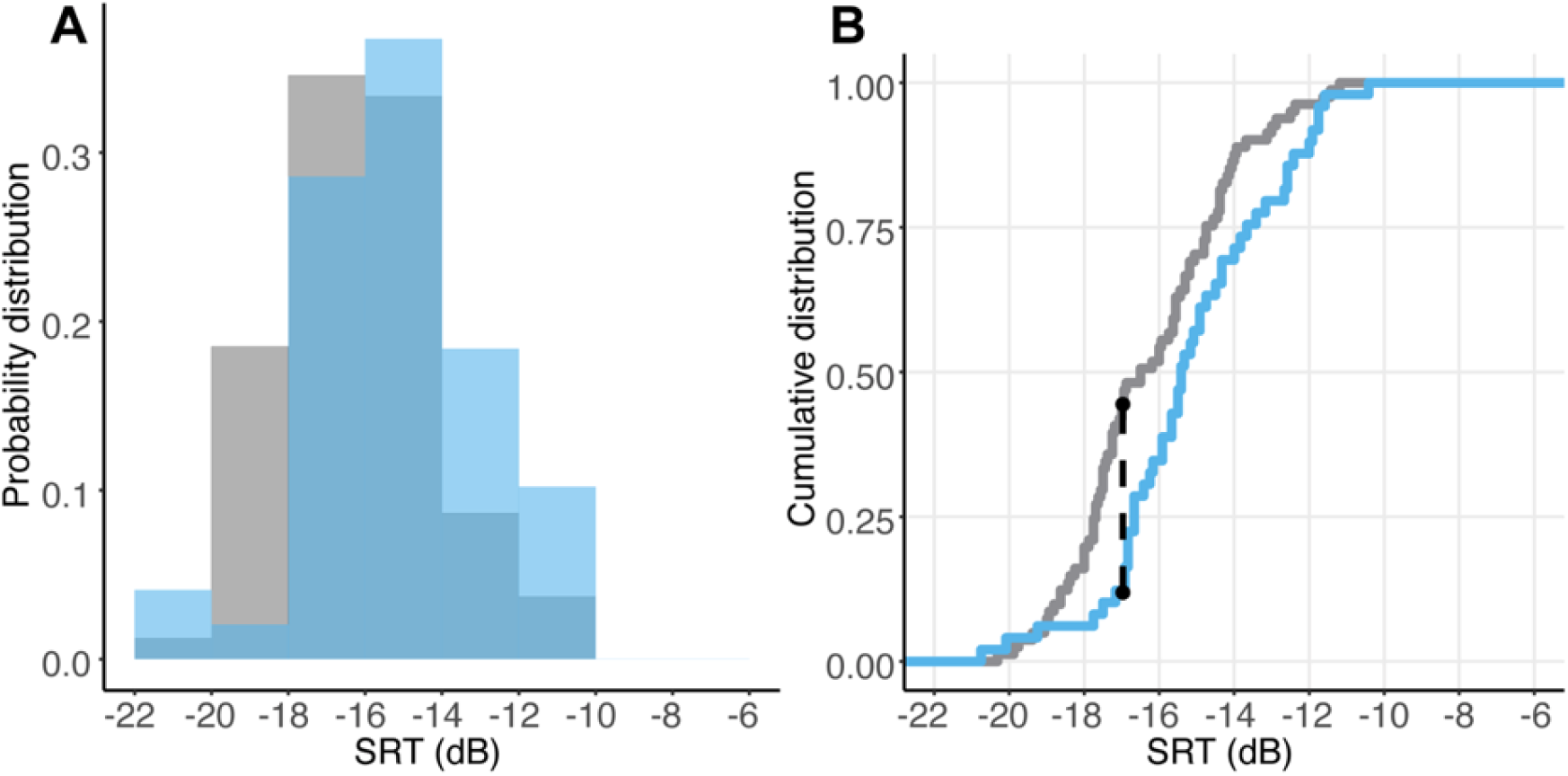
**A.** Probability density distributions of the in-lab (grey) and online (blue) groups. **B.** Cumulative distribution of the in-lab and online groups. The black dotted line indicates the SRT at which the greatest distance between the two distributions was observed. Overall the data pattern is consistent with a right-ward shift (towards higher SRTs) of the online distribution.

The overall pattern of results demonstrates that, relative to the in-lab cohort, fewer people in the online group achieved very low (< −17dB) thresholds, suggesting that online testing may provide a less accurate measure of listeners’ speech-in-noise detection performance. The differences between the online and in-lab groups may arise due to a poorer control of participants’ listening environment and / or motivation.

## Experiment 2

### Methods

#### Participants

Two hundred young, normal-hearing listeners ranging in age from 22 to 30 years (128 females, mean age 26 ± 2.5) were recruited online as described in Experiment 1. They were randomly assigned to one of two experimental groups. One group (N = 100, 62 females) received a performance-based monetary bonus (up to £ 5) on top of the base-pay **(BONUS+).** The other group (N = 100, 66 females) received no bonus **(BONUS-).**

#### Stimuli and procedure

The procedure was similar to that described for Experiment 1. The BONUS+ and BONUS- groups received identical instructions and feedback. Encouraging language was used to maximize participant motivation. After each run, the achieved threshold was displayed and participants were challenged to try to ‘beat their score’ in the next run. The BONUS+ group were additionally informed that each threshold was associated with a monetary bonus. They were told that at the end of the experiment they would receive the bonus (up to £ 5) associated with the best threshold reached (for example, if they reached thresholds −10, −15, −17, −10 over the runs, they were payed a bonus associated with threshold −17). At the end of each run, participants were shown the current threshold and the associated bonus, but also the bonus they could receive if they improve their threshold in the following run. The bonus was preassigned to SNR values from −1 to −28 (in steps of 1) through an exponential function, so that improvements at lower, more difficult thresholds were rewarded more than improvements at levels expected to be easily reached by young normal-hearing listeners. As in Experiment 1, following the main task participants answered a set of questions about their listening environment. They were also asked to answer on a scale from 0 to 10 (0 = not at all, 10 = a lot) how motivated they were in performing the task, and how engaging they found the task to be.

The base-pay was set to £ 2 (for 15 minutes). The average obtainable bonus was £ 2 (range £ 0-5) therefore allowing participants to double their pay. To avoid bias in the selection process, participants were unaware of the possibility to be assigned to a bonus group when they signed up to the study. The BONUS+ group were only informed of the bonus at the instructions stage.

## Results

Both the BONUS+ and BONUS- groups reported a similar level of environmental noise (BONUS+ = 1.87 ± 2.75; BONUS- = 1.87 ± 2.75; t-test: t(2,198) = 1.203, p = .230). However, to focus on the effect of bonus on performance, we excluded those participants who reported a level of noise >= 5 (on a scale from 0-10) resulting in the exclusion of ~15 participants from each group (final numbers: BONUS+ N = 84; BONUS- N = 90). Figure 2 shows the probability density function (panel A) and the cumulative distribution function (panel B) of the SRT obtained for the BONUS+ (mean SRT = −16.1 dB, SD = 2.54), and BONUS- (mean SRT = −15.1 dB, SD = 2.33) groups. Data from the in-lab group (see Experiment 1) are also provided as a benchmark.

**Figure 2.**
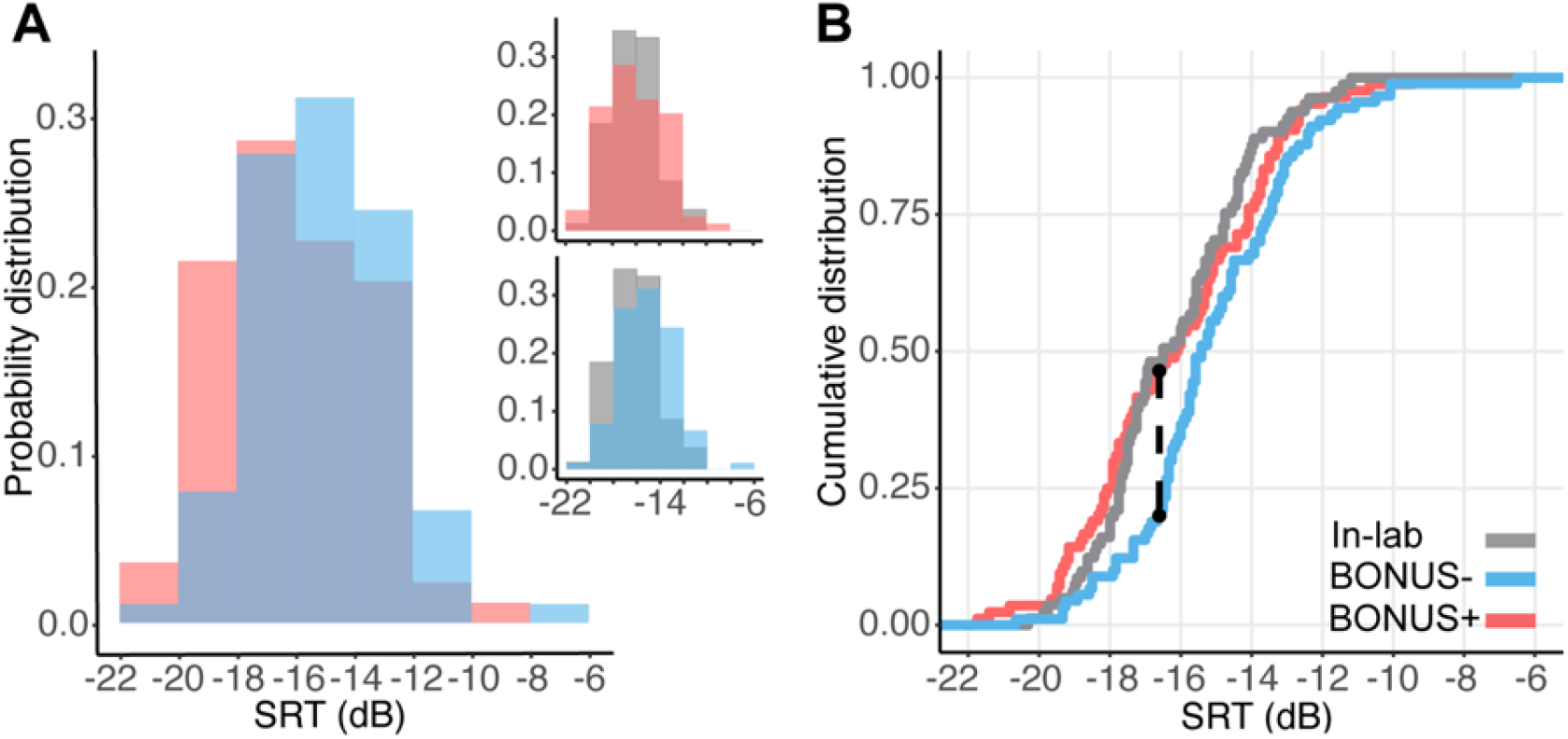
A. Probability density distributions (relative proportion) of the online BONUS+ (pink) vs. the BONUS- (blue) groups. The insets show the probability density distributions of the BONUS+ (top) and the BONUS- (bottom) groups against the in-lab sample. B. Cumulative distribution of the BONUS+ and BONUS- groups. The data from the in-lab (grey) are plotted as benchmark. The black dotted line indicates the SRT at which the greatest distance between the BONUS+ and BONUS- distributions is observed. Overall the data pattern is consistent with a left-ward shift (towards lower SRTs) of the BONUS+ relative to the BONUS- groups.

KS tests indicated a significant difference between the BONUS+ vs BONUS-distributions (D = .276, p = .003), revealing better performance in the BONUS+ compared with the BONUS- group. The maximum difference occurred at −16.6 dB, which was reached by 47% of the BONUS+ and only by the 21% of the BONUS- group. The comparison of these two distributions with the in-lab distribution indeed showed that the BONUS+ performance was similar to the in-lab one (D = .127, p = .519), whilst the BONUS- was different (D = .304, p = .001). The results thus indicate that the provision of a bonus increased the proportion of high performing participants in the online group to the levels exhibited by the in-lab cohort.

In an additional analysis we compared the in-lab group with the online data pooled from the online group of Experiment 1 and the BONUS- group of Experiment 2 (for a total of N = 132, excluding participants who reported a level of background noise >= 5; note all results hold even without excluding participants based on noise reports). A KS test confirmed that online performance in the absence of an additional bonus is worse than that obtained in-lab (D = .318, p < .001), in line with what was observed in Experiment 1.

Consistent with the interpretation that offering a bonus increased motivation, the BONUS+ group reported higher ratings of task engagement [BONUS+ = 9.1 ± 1.2; BONUS- = 8.4 ± 1.5; from range of 0-10; t(2,98) = 2.68, p = .009] and motivation [BONUS+ = 9.2 ± 1.12; BONUS- = 8.4 ± 1.3; from range of 0-10; t(2,98) = 3.212, p = .002] compared with the BONUS- group.

## Discussion

We report two main findings: Firstly, we showed that the SRT of blindly recruited online participants was poorer than that observed among an age-matched control group in the lab setting. Second, we demonstrated that the provision of a small performance-based monetary bonus improved online listeners’ speech-in-noise performance to levels similar to those observed in the lab setting.

The results from Experiment 1 revealed that the distribution of the SRT in the online group differed from that obtained from the in-lab cohort: In the lab, 47% of listeners achieved an SRT below ~ −17 dB. In contrast, within the online cohort only 12% of participants reached that threshold. This discrepancy is relevant to consider when using remote testing to build normative data, or to accurately estimate hearing loss across the population.

In Experiment 2, we showed that a performance-based monetary bonus increased the proportion of highly performing participants (SRT < −17 dB) up to levels similar to those observed in the lab. This suggests that the difference in performance between the online and in-lab groups observed in Experiment 1 is not mainly driven by constraints to the sound environment but rather associated with reduced task-engagement among the online participants.

With the blooming of online experiments, it is important to understand how we can improve the quality of data obtained in remote auditory assessments (Leensen, De Laat, Snik, & Dreschler, 2011; Milne et al., 2020; Slote & Strand, 2016). Our finding that reward increased the proportion of participants who achieved low SRTs demonstrates that participant attention, motivation and commitment are important factors to consider when auditory tests involving effortful listening are conducted online. Higher task-engagement in the in-lab than in the online population probably result from several factors that characterise the laboratory experience: the authority of experimenter, the absence of temptation/distractions, the effort taken to come to the lab, etc. All these factors are likely to make in-lab participants already quite motivated. Similar considerations may apply to certain online testing situations. For example, participants in remote clinical assessments are likely to be highly intrinsically motivated to do their best, as revealed by studies reporting similar results between testing in the clinic and at home (de Graaff, Huysmans, Merkus, Theo Goverts, & Smits, 2018; Whitton, Hancock, Shannon, & Polley, 2016). However in many cases, online participants are unsupervised and anonymous, and often mainly motivated by financial incentives (Buhrmester, Kwang, & Gosling, 2011; Litman, Robinson, & Rosenzweig, 2015). In a recent in-house survey conducted by Prolific.co approximately 50% of the surveyed users stated that the amount of pay is the factor that most motivates them to take part in a study (https://prolific2.typeform.com/report/PoUZHEmk/ttebnlTEllbRvdcg). Therefore, an efficient method for increasing task-engagement is a monetary bonus. This consideration is also supported by the fact that the BONUS+ group in Experiment 2 reported higher ratings of task engagement and motivation compared with the BONUS- group.

Previous studies suggest that performance on many crowdsourcing tasks does not differ, and sometimes even exceeds that measured in the lab (Gallun et al., 2018; Hauser & Schwarz, 2016; Lelo de Larrea-Mancera et al., 2020; but see Harrison & Müllensiefen, 2018; Slote & Strand, 2016). Additionally, monetary incentive (amount of pay) does not necessarily affect performance (van den Berg, Zou, & Ma, 2019): For example, previous studies reported no modulatory effects of amount of monetary incentive on the quality of online performance in tasks such as speech transcriptions (Marge, Banerjee, & Rudnicky, 2010). Internal consistency in psychological surveys and attention in following instructions were also unaffected by different levels of payment (Buhrmester et al., 2011; but see Litman et al., 2015). However, the impact of incentive may depend on the kind of task under investigation. Financial incentives may have little effect on performance when the task is too easy or when return on effort is low, e.g. when it is very hard to improve performance (Camerer & Hogarth, 1999). Our finding that reward influences performance in the CCRM task is possibly linked to the fact that the return on effort is high: the task relies on attention to fine perceptual details and increasing effort has the potential to lead to a notable improvement in performance.

The effect of incentives on performance may also be nuanced by how the reward is delivered. Shen & Chun (2011) demonstrated that reward can encourage participants to perform better when it is progressively increased from trial to trial, but not when the same high reward level is maintained. Furthermore, the effect of reward in Shen & Chun (2011) appeared to hold true even when the ultimate reward was success in a competition (e.g., a monetary reward assigned to the top 10% participants based on performance), rather than money itself (e.g., performance-based earning with no competition). Therefore, particularly in experiments that require many trials, a competitive setting may be a more effective incentive than a small (a few cents) reward per trial.

## Conclusions

How reward might motivate performance is an empirical question and a longstanding object of debate. Accumulating evidence suggests that reward does seem to matter particularly in tasks where performance depends on effortful engagement (Camerer & Hogarth, 1999). The CCRM task used here is analogous to many threshold-based tasks commonly used in auditory research. The observed effect of bonus on performance should thus generalize to other auditory tasks, helping to motivate participants to exert the extra bit of effort that is needed when the task becomes just-doable.

## Open practice Statements

Stimuli and code implementing the CCRM test can be found in Gorilla Open Materials: https://gorilla.sc/openmaterials/171870.

